# Transcranial Near-Infrared Light: Dose-Dependent Effects on EEG Oscillations but not Cerebral Blood Flow

**DOI:** 10.1101/837591

**Authors:** Vincenza Spera, Tatiana Sitnikova, Meredith J. Ward, Parya Farzam, Jeremy Hughes, Eric Bui, Marco Maiello, Luis De Taboada, Michael R. Hamblin, Maria Angela Franceschini, Paolo Cassano

## Abstract

**Objective:** Our objective was to assess whether transcranial photobiomodulation (tPBM) with near-infrared light (NIR) shows modulatory effects on cerebral electrical activity through electroencephalogram (EEG) and cerebral blood flow (CBF).

**Background:** tPBM has emerged as a novel intervention for several neuropsychiatric conditions due to its neuroprotective and neuroenhancement effects.

**Methods:** We conducted a single-blind, sham-controlled pilot study to test the effect of continuous (c-tPBM), pulse (p-tPBM) and sham (s-tPBM) transcranial photobiomodulation on EEG oscillations and CBT using diffuse correlation spectroscopy (DCS) in a sample of ten healthy subjects [6 F/ 4 M; mean age 28.6 ± 12.9 (SD) years]. c-tPBM NIR (830 nm; 54.8 mW/cm²; 65.8 J/cm²; 2.3 kJ) and p-tPBM (830 nm; 10Hz; 54.8 mW/cm²; 33%; 21.7 J/cm²; 0.8 kJ) were delivered concurrently to the frontal areas (F4, Fp2, Fp1, F3 - total surface 35.8 cm²) by four LED clusters (TPBM-1000 Litecure). EEG and DCS recordings were performed weekly before, during, and after each tPBM session (in sequence: c-tPBM, s-tPBM and p-tPBM).

**Results:** Only c-tPBM significantly boosted gamma (t = 3.02, df = 7, p < .02) and beta (t = 2.91, df = 7, p < .03) EEG spectral powers in eyes-open recordings and gamma power (t = 3.61, df = 6, p < .015) in eyes-closed recordings, with the largest effects in the posterior regions. There was no significant effect of NIR-tPBM on CBF compared to sham.

**Conclusions:** Our data suggest a dose-dependent effect of tPBM with NIR (c-tPBM) on cerebral gamma and beta neuronal activity. Altogether, our findings support the neuromodulatory effect of transcranial NIR.

## 1. Introduction

Transcranial Photobiomodulation (tPBM) with Near-infrared radiation (NIR) is a novel intervention based on the use of low-level lasers or light-emitting diodes (LEDs) that has recently emerged as a potential valuable therapy for a range of neuro-psychiatric conditions. tPBM may have therapeutic effects in subjects with stroke, traumatic brain injury, neurodegenerative disorders and major depressive disorder (Cassano *et al*., 2018), as well as pro-cognitive benefits in healthy populations (Barrett & Gonzalez-Lima, 2013; Blanco, Maddox, & Gonzalez-Lima, 2017; Hamblin, 2018). Recent findings have pointed out the beneficial effect of tPBM in the augmentation of cognitive functions, such as memory and attention, in addition to emotional functions. In a placebo-controlled randomized study in healthy volunteers, Barrett and Gonzalez-Lima (Barrett & Gonzalez-Lima, 2013) showed a significant improvement in performance on the frontal lobe cognitive tasks and in the emotional state after tPBM (1064 nm laser, 250 mW/cm², 60 J/cm², 13.6 cm² x 2 sites) delivered with continuous light to participants’ forehead.

The mechanisms through which tPBM with NIR or red light, delivered to the scalp of patients, may influence the adjacent cortical areas of the brain are poorly understood. One promising hypothesis is that tPBM may influence brain energy metabolism through promoting the mitochondrial function. The primary source of intracellular energy, adenosine triphosphate (ATP), which is critical to sustaining neural activity, is largely produced in mitochondria through the process of oxidative phosphorylation. This process involves a respiratory chain of five enzyme complexes which, if altered, would influence ATP synthesis. It has been suggested that tPBM, by delivering photons (energy particles) to the tissue, may promote one of such complexes – cytochrome c oxidase (CCO) or respiratory chain complex IV –restoring or enhancing ATP production and leading to more energy available for neuronal activity (Hennessy & Hamblin, 2017). Several studies reported an upregulation effect on CCO from both the LED and laser light therapy at or close to 830 nm, which led to the neuronal increase in energy production (Mochizuki-Oda *et al*., 2002; Wong-Riley *et al*., 2005). Mitochondria dysfunction has been suggested in many common neuropsychiatric disorders (Marazziti *et al*., 2012), and tPBM may enable compensation for such dysfunction, to restore cognitive capacity.

More active mitochondria would support higher oxygen/glucose consumption, which might stimulate cerebral blood flow (CBF) to deliver such nutrients. Intriguingly, a preclinical study by Uozumi *et al*. (2010) reported an increase of 30% of CBF after transcranial NIR laser irradiation at 808 nm wavelength for 45 min (at a power density of 1.6 W/cm²), not related to heating (Uozumi *et al*., 2010). In healthy human participants and clinical patients, cerebral oxygenation and cerebral blood flow were also found to increase (Tian, Hase, Gonzalez-Lima, & Liu, 2016) (Salgado, Zangaro, Parreira, & Kerppers, 2015). Through its effects on both cerebral blood flow and brain metabolism (Blanco *et al*., 2017), tPBM is considered a potential non-invasive therapy for cognitive impairment based on both animal and human studies (de la Torre, 2016).

More active mitochondria have also been demonstrated to influence brain activity, especially oscillations at high frequencies. Extensive research in slice cultures of hippocampus has documented the link between the mitochondrial function and fast neural oscillations in the gamma band (∼30-90Hz) (Galow *et al*., 2014; Whittaker, Turnbull, Whittington, & Cunningham, 2011). Gamma frequency activity is believed to arise from the interplay between cortical inhibitory interneurons and excitatory principal neurons – the mechanism being: high rates of interneuron firing are required to synchronize function of principal neurons (Kann, Papageorgiou, & Draguhn, 2014) (Buzsaki & Wang, 2012). The high metabolic demands of such interneuron activity may be linked to the activity of the mitochondrial respiratory chain in these cells. Intriguingly, the parvalbumin-positive interneurons, which are critical to gamma oscillations, show higher levels of mitochondrial CCO when compared with principal neurons (Gulyas, Buzsaki, Freund, & Hirase, 2006). Thus, by stimulating the mitochondrial enzyme, CCO, tPBM may deliver targeted stimulation of the parvalbumin-positive interneurons and would be expected to enhance gamma oscillations. Gamma oscillations provide a fundamental mechanism of complex neuronal information processing in the hippocampus and neocortex of mammals. This type of brain electrophysiological activity has been implicated in higher brain functions such as sensory perception (Gross, Schnitzler, Timmermann, & Ploner, 2007), motor activity (Nowak, Zich, & Stagg, 2018) and memory formation (Sederberg *et al*., 2007), and are impaired in many common neuropsychiatric disorders (Herrmann & Demiralp, 2005). However, only limited evidence is currently available on the effects of tPBM with NIR on the high-frequency neural oscillations in the human brain and on its ideal dosimetry.

The goal of the current study was to elucidate the physiological processes that may underlie the pro-cognitive effects of tPBM with NIR by tracking the effects of two different treatment doses on simultaneous recordings of neural activity oscillations and CBF. We delivered 830-nm tPBM by a four LED clusters device to the forehead of healthy participants and employed electroencephalography (EEG) to quantify electrophysiological brain oscillations and diffuse correlation spectroscopy (DCS) to index CBF. We anticipated that tPBM would stimulate the CBF and the electrophysiological oscillations, especially in the high frequency bands. We also expected that pulsed light (p-tPBM) would exert a stronger effect compared to continuous light therapy (c-tPBM), which would be consistent with prior reports (Lapchak & De Taboada, 2010) (Brondon, Stadler, & Lanzafame, 2009) (Lapchak, Salgado, Chao, & Zivin, 2007). Secondary aims were to assess the safety and tolerability of NIR-light therapy delivered transcranially.

## 2. Materials and methods

This single-site study –*Transcranial Near-Infrared Light in Healthy Subjects: a Cerebral Blood Flow Study with Diffuse Correlation Spectroscopy (NIR-flow)* – was approved by the Massachusetts General Hospital (MGH) institutional review board (IRB). The main sources of recruitment were email and website ads through the *Partners Health Care* internal portal for clinical trials. The study clinicaltrials.gov identifier was NCT03740152.

### 2.1 Inclusion and exclusion criteria

Subjects (age 18-70) eligible for study participation were healthy by Structured Clinical Interview for Diagnostic Statistical Manual-IV (SCID) criteria. Subjects were enrolled in the study after providing written informed consent. Female subjects of childbearing potential needed to consent (without any element of coercion) to use a double-barrier method for birth control if sexually active; pregnancy and lactation were exclusionary. Other exclusionary criteria included any current psychiatric disorder, substance or alcohol use disorders (prior 6 months), lifetime psychotic episodes, bipolar disorder, unstable medical or neurological illness, recent history of stroke, active suicidal and homicidal ideation. In addition, the following criteria, potentially influencing light penetration and safety, were also exclusionary: any use of light activating drugs (prior 14 days), having a forehead skin condition (such as tattoo or open wound) and having a head-implant. Out of eleven eligible subjects, ten subjects were followed for the entire five-week study period and completed a sequence of three sessions of tPBM and sham. The study involved screening (week 1), 3 tPBM sessions (weeks 2-4), and one follow-up visit (week 5).

### 2.2 Study design, blinding and assessment schedule

The study included three sequential sessions, separated by at least one week, which entailed three different modes of operation of the tPBM device (Figure 1). Participants received one session of continuous light treatment (c-tPBM), followed by a session with sham treatment (s-tPBM), which was followed by a session with pulse light treatment (p-tPBM). While the sequence (c- tPBM, s-tPBM, p-tPBM) was the same for all subjects, the subjects were blind to the specific mode of device operation used in each session. Simultaneous recordings of EEG and DCS were conducted as subjects rested (with no cognitive task) before, during, and after each tPBM session. EEG was also recorded while participants performed a working memory (2-back) task: once at baseline –before the c-tPBM session– and after each tPBM session. NIR light was administered by a four LED clusters device (LiteCure® TPBM-1000) which had the flexibility to deliver either continuous light (c-tPBM), pulse light (p-tPBM), or sham. In both c-tPBM and p- tPBM mode the TPBM-1000 device delivered therapeutic NIR energy. In sham (s-tPBM) the device didn’t deliver any light energy. The apparent behavior (i.e. the performance/output of all visible and audible indicators) of the device in any of the three programmed treatment modalities was identical. Because of the low average irradiances delivered in both c-tPBM and p-tPBM modes and because of heat sinks incorporated in the device, the subjects did not experience any skin warming from NIR. These features of the design ensured that the study was single-blind. The device was designed to shut off automatically if skin warming above 41C was detected. Tolerability was assessed through clinician inquiry and through a self-report scale (SAFTEE-SI), which were administered before the first session (c-tPBM), as a baseline measure, and one week after each session (c-tPBM, s-tPBM, p-tPBM) to examine treatment-emergent side effects.

**Figure 1.**
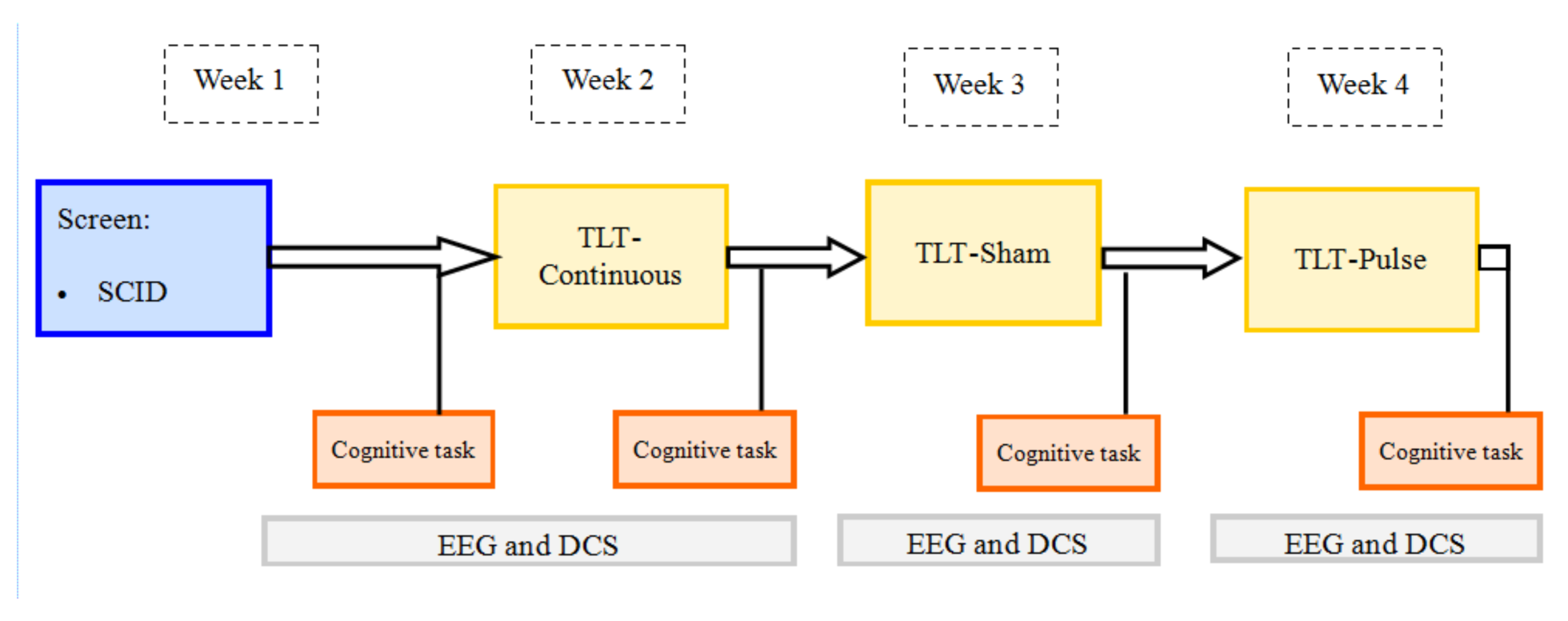
Flowchart of the study procedures

### 2.3 Intervention parameters (tPBM)

NIR light at 830 nm was delivered with the TPBM-1000 device (Figure 2) to four EEG sites (Fp1, Fp2, F3, F4) targeting frontal poles and dlPFC, covering a total surface active treatment area of 35.8 cm^2^ [(11.52 cm^2^x2; Fp1, Fp2) + (6.38 cm^2^x2; F3, F4)] with an irradiance of 54.8 mW/cm^2^ (average in c-tPBM and peak in p-tPBM), an average fluence of 65.8 J/cm^2^ in c-tPBM and of 21.7 J/cm² in p-tPBM, and a total energy of 2.3 kJ in c-tPBM and 0.8 kJ in p-tPBM (Table 1). The additional parameters used for pulsed light were the following: frequency 10 Hz, duty cycle 33%. The duration of irradiation was 20 minutes (the 4 sites were irradiated concurrently). The entire sessions lasted about 1.5 hours. The additional time was needed to have the subject complete a urine toxicology test and self-report forms, to prepare the subject, to place the necessary protections (e.g. goggles), to inspect the subject’s skin, to complete DCS and EEG (before, during and after tPBM), to set the tPBM devices, and to give the subject time to rest after the irradiation. tPBM was administered by licensed physicians (i.e. MDs) who were on study staff and trained to for the use of TPBM-1000.

**Figure 2.**
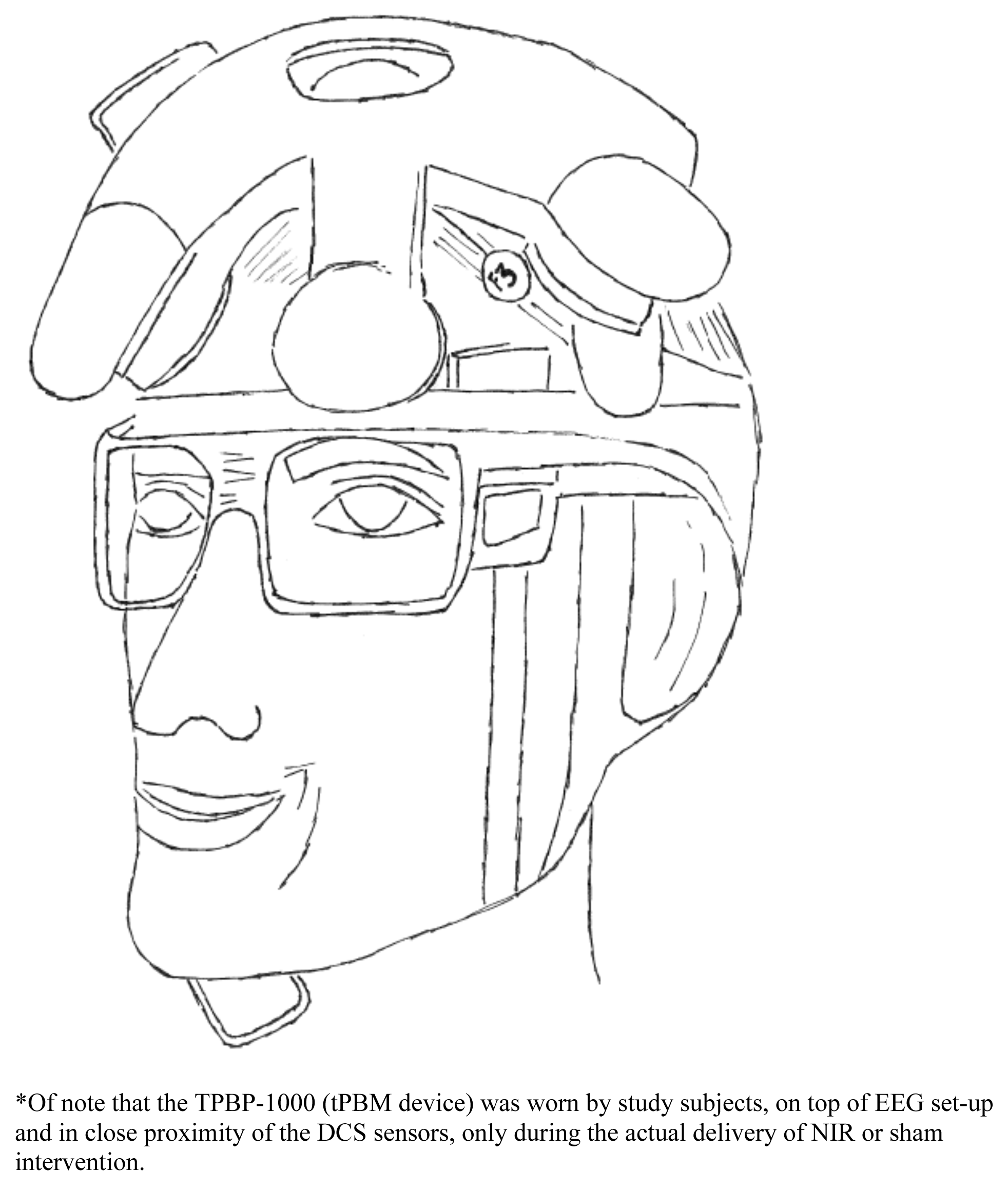
Set up of TPBM-1000, DCS and EEG devices, mounted on study subject’s head*

**Table 1:** Treatment parameters for tPBM delivered as continuous wave (c-tPBM) and as pulse wave (p-tPBM)

|  | **c-tPBM** | **p-tPBM** |
| --- | --- | --- |
| Wavelength | 830 nm | 830 nm |
| Irradiance (peak) | 54.8 mW/cm<sup>2</sup> | 54.8 mW/cm<sup>2</sup> |
| Pulsing rate | - | 10 Hz |
| Duty cycle | 100% | 33% |
| Average Power | ~2W | ~0.7W |
| Peak Power | ~2W | ~2W |
| Fluence | 65.8 J/cm<sup>2</sup> | 21.7 J/cm<sup>2</sup> |
| Duration of t-PBM session | 20 min | 20 min |
| Treatment window | 35.8 cm<sup>2</sup> | 35.8 cm<sup>2</sup> |
| Cumulative dose | 2.3 kJ | 0.8 kJ |

### 2.4 Electroencephalographic activity (EEG)

Electroencephalography (EEG) is an imaging technique that can track electrophysiological activity of the brain with a high temporal resolution. Previously, EEG has been used to evaluate neurophysiological brain activity both during resting ‘task free’ scans –to study patterns of spontaneous dynamics of brain activity (Rosazza & Minati, 2011) and during task performance to examine how brain activity is modulated by the demands of a task (Bridwell *et al*., 2018). Fluctuations in the EEG signal have been studied in several frequency bands, including delta (0.5–3.5 Hz), theta (4–8 Hz), alpha (8–12 Hz), beta (12–30 Hz) and gamma (>30 Hz). Such patterns of slow and fast fluctuations have been linked to different states of mental activity, and different sensory and cognitive processes (Herrmann, Struber, Helfrich, & Engel, 2016).

The Stat® X24 wireless EEG system (Advanced Brain Monitoring, Carlsbad, CA) was used for EEG data acquisition. The Stat X24 combines battery-powered hardware with a preconfigured sensor strip for recording twenty channel monopolar EEG. The Stat X24 provides 19 EEG channels in accordance with the International 10-20 system, including Fz, F3, F4, Cz, C3, C4, P3, P4, Pz, O1, O2, T5, T3, F7, Fp1, Fp2, F8, T4, and T6, plus adds POz. EEG (sampling rate, 256 Hz, band pass filter: 0.1 Hz high-pass, 100 Hz fifth order low-pass, referenced to linked mastoids) was recorded and analyzed during rest before and after each tPBM session and during each 2-back task session. Additionally, electrocardiogram (ECG) was recorded from collarbone left/right locations to enable removal of cardio-artifacts. Independent component analysis (ICA) was used to remove any artifact due to eye-blinks (using a software package from OHBA: https://github.com/OHBA-analysis/osl-core). During each resting state recording, participants reclined in a comfortable chair for 5 minutes with eyes open (while looking at a cross shown on a computer screen) and 5 minutes with eyes closed. Participants were asked to relax and avoid any eyebrow movements or clenching their jaw. Because the state of brain function was expected to change from before to after tPBM not only due to the light treatment itself, but also due to relaxing in a darkened room for a considerable time during treatment (Ward *et al*., 2013), our primary comparison of interest was between EEG power spectral density (PSD) in post and pre tPBM recordings, after subtracting corresponding PSD recorded on a sham treatment visit. Because we hypothesized that tPBM would have an effect on the brain activity in higher frequencies, we limited our analyses to the gamma, beta, alpha bands.

### 2.5 Cerebral blood flow (DCS)

Diffuse correlation spectroscopy (DCS) is an optical technique that uses the temporal fluctuations of near-infrared (NIR) light to measure cerebral blood flow (CBF) directly and non-invasively. DCS devices to date are not under Food and Drug Administration (FDA) investigational device exemption (IDE) regulations, since they are considered non-significant risk. A homebuilt DCS system was used with a long-coherence length 852-nm laser, 4 photon counting avalanche photodiode detectors, and a custom FPGA correlator. The DCS system was placed adjacent to one of the tPBM sources, at either the left or right frontal pole. The DCS light source power at the optical probe was below 40 mW, and the beam diameter was >1 mm (to comply with ANSI standards for skin exposure). The detectors were arranged at different distances from the source. In particular, we had one detector at 5 mm separation from the source to detect superficial blood flow changes and 3 detectors at 2.5 cm to detect blood flow in deeper tissue. The DCS measurements were done before and after tPBM and every 1 min for 10 sec during tPBM, by turning off the tPBM light and turning on DCS source and detectors (to avoid detectors damage by the powerful tPBM light). Blood flow index at the short separation (BFi) was calculated for each detector by assuming a fixed absorption and scattering across subjects. CBFi at the three large separations was averaged together to reduce noise. Temporal changes of CBFi (large separation), of BFi (short separation), and the subtraction of the two were considered for the statistical analyses.

### 2.6 Cognitive Task (2-Back)

In the 2-back task, participants were presented with a sequence of English letters, and were asked to press a button when the current letter matched the one from 2 steps earlier in the sequence. Such 2-back task is believed to capture the engagement of working memory – one of the cognitive functions supported by the dorsolateral prefrontal cortex (dlPFC). To perform this task, it is not enough to simply keep a representation of recently presented items in short-term memory; the working memory buffer also needs to be updated continuously to keep track of what the current stimulus must be compared to. In other words, the participant needs to both maintain and manipulate information in working memory (Chai, Abd Hamid, & Abdullah, 2018). Our primary interest was in whether we could detect differences in the task performance after any of the active tPBM sessions (c-tPBM and p-tPBM), relative to the performance after a sham (s-tPBM). Accordingly, participants were asked to perform 2-back task after each active/sham session. To reduce burden on the participants, we administered the 2-back task before tPBM only at the first visit (c-tPBM). We decided to present only results comparing n-backs pre- and post-c-tPBM, based on the DCS and EEG results across the three sessions. To examine how well each participant was able to perform the 2-back task, we computed both the accuracy in detecting the target letters –2-back hits and the number of false positive responses– and reaction times.

### 2.7 Statistical analysis

Paired t-tests were performed to compare the effect of different light treatments on both EEG and CBF/DCS recordings. Due to the pilot nature of this study, since we recruited a small number of participants, all statistics are reported at .05 level of significance (no correction for multiple comparisons).

## 3. Results

Table 2 describes the demographic characteristics of the sample. The ten subjects (six females/ four males) had a mean age of 28.6 ± 12.9 (SD). The tPBM sessions were well tolerated without any serious adverse event. As measured by the SAFTEE scale, administered one week after each session (c-tPBM, s-tPBM, p-tPBM), three subjects developed one or more adverse events: one subject (#1) experienced ‘feeling drowsy / sleepy’ and ‘weakness / fatigue’, one subject (#5) experienced ‘trouble concentrating’ and one subject (#8) reported ‘blurred vision’ and ‘nausea/vomiting’. Of note, our study included multimodal integration of several technologies and techniques –including EEG and DCS recording, tPBM delivery, 2-back cognitive and self-report assessments– which presented several logistical challenges. This complex design –while offering new, multimodal insights on the neurophysiology of tPBM– led to the loss of data in some recording sessions, due to insufficient quality or to participants’ time constraints. We report the number of included datapoints for each comparison below.

**Table 2:** Demographic characteristics on healthy subjects enrolled in the study

| **Subject** | **Age** | **Gender** | **Ethnicity** | **Race** | **Skin color** |
| --- | --- | --- | --- | --- | --- |
| **#1** | 56 | F | Hispanic | White | 5 out of 10 |
| **#2** | 28 | F | Not Hispanic | White | 2 out of 10 |
| **#3** | 49 | M | Not Hispanic | White | 1 out of 10 |
| **#4** | 23 | F | Not Hispanic | Asian | 4 out of 10 |
| **#5** | 23 | F | Not Hispanic | White | 2 out of 10 |
| **#6** | 24 | M | Not Hispanic | White | 1 out of 10 |
| **#7** | 20 | F | Not Hispanic | Asian | 2 out of 10 |
| **#8** | 21 | F | Hispanic | More than one race | 4 out of 10 |
| **#9** | 22 | M | Not Hispanic | Asian | 2 out of 10 |
| **#10** | 20 | M | Hispanic | White | 1 out of 10 |

### 3.1 Electroencephalographic activity (EEG)

#### Open Eyes EEG Recordings at Rest

We observed that continuous light treatment (c-tPBM), but not pulsed light treatment (p- tPBM), influenced EEG power spectrum density (PSD). Figure 3 shows the results for resting state recordings with eyes open. Results for c-tPBM (CW) treatment (n = 8) are shown in the left panel. We computed pre-light-treatment (pre-tPBM) PSD by subtracting pre-treatment recording at the sham session from pre-treatment recording at the c-tPBM session. Similarly, we computed post-light-treatment (post-tPBM) PSD by subtracting post-treatment recording at the sham session from post-treatment recording at the c-tPBM session. These PSD metrics gave us the change value that was likely due to c-tPBM treatment, while controlling for the sham effect of merely staying in a dark room with a light-treatment device applied but not active. The scalp maps on top of Figure 3 show PSD data averaged across participants for alpha (8-12 Hz), beta (12-30Hz), and lower gamma (30-55Hz) EEG recordings, with pre-tPBM PSD data in the bottom row and post-tPBM data in the top row. There is a widespread increase in the activity PSD over frontal-central scalp regions post-tPBM (post-LT), relative to pre-tPBM (pre-LT), especially in the gamma band. The spectrogram for this data, averaged across all EEG channels, is shown below Figure 3, the solid lines show group average and the shaded areas shows the standard error of the mean across participants. The global PSD increase in the post-tPBM relative to pre-tPBM recordings is larger for higher frequencies. This global effect reached significance for gamma (mean change in γ-PSD_[POST (CW - Sham) – PRE (CW - Sham)]_ .2426 ± .2276 SD, t = 3.02, df = 7, p < .02) and beta (mean change in β-PSD_[POST (CW - Sham) – PRE (CW - Sham)]_ .1391 ± 1354 SD, t = 2.91, df = 7, p < .03) bands, but it was not significant in the alpha band (mean change in α-PSD_[POST (CW - Sham) – PRE (CW - Sham)]_ .0822 ± .1665 SD, t = 1.40, p > .2). Results for p-tPBM (PW) treatment (n = 9) are shown in the right panel. Again, we computed pre-tPBM PSD and post-tPBM PSD by subtracting pre- and post-treatment recordings at the sham session from pre- and post-tPBM recordings at the p-tPBM session, respectively. Even though there appears to be a frontal increase in PSD due to p-tPBM treatment, the result did not reach significance.

**Figure 3.**
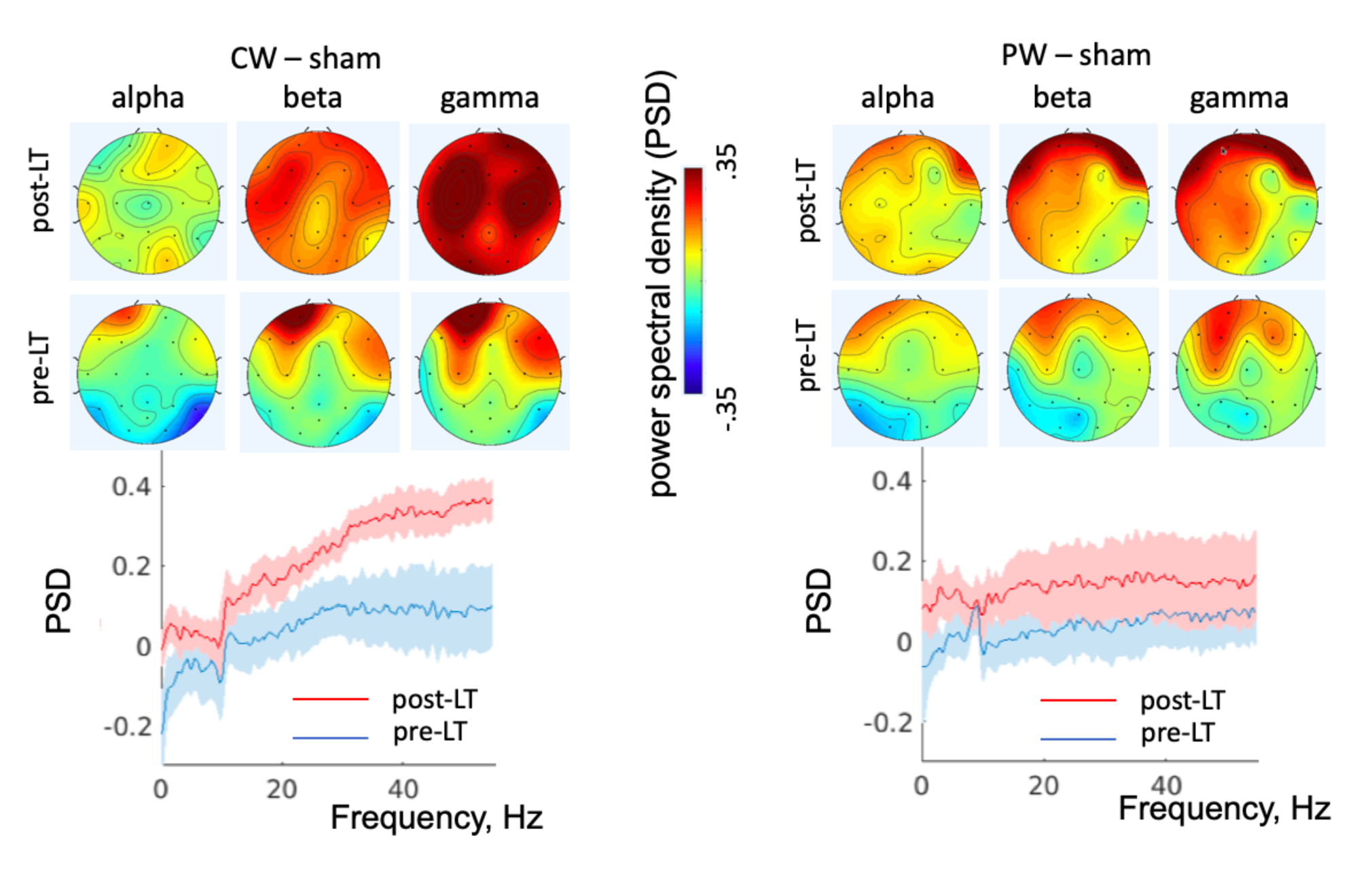
Eyes-open resting state EEG: spectrogram differences before and after tPBM with continuous (CW) and pulse (PW) light (LT).

#### Closed Eyes EEG Recordings at Rest

Figure 4 shows the results for resting state recordings with closed eyes, which were obtained according to the same procedure as with open eyes. Results for c-tPBM (CW) treatment (n = 7) are shown in the left panel. Again, there was a widespread increase in the activity power in post-tPBM relative to pre-tPBM scans for gamma and beta bands. In the alpha band, activity power was high both pre-tPBM and post-tPBM, which would be expected as alpha is known to increase in power when participants close their eyes. The global effect is illustrated in the lower panel of the Figure. As with global PSD increases in *Open Eyes EEG Recordings at Rest*, the PSD increased with EEG frequency. The effect reached significance in the gamma band (mean change in γ-PSD_[POST (CW - Sham) – PRE (CW - Sham)]_ .3654 ± .2678 SD, t = 3.61, df = 6, p < .015), but was not significant in the beta (mean change in β-PSD_[POST (CW - Sham) – PRE (CW - Sham)]_ .1377 ±.2069 SD, t = 1.77, df = 6, p > .1) or alpha (mean change in α-PSD_[POST (CW - Sham) – PRE (CW - Sham)]_ -.0223 ± .2233 SD, t = .026, df = 6, p > .8) bands. Results for p-tPBM (PW) treatment (n = 8) are shown in right panel. None of the result reached significance.

**Figure 4.**
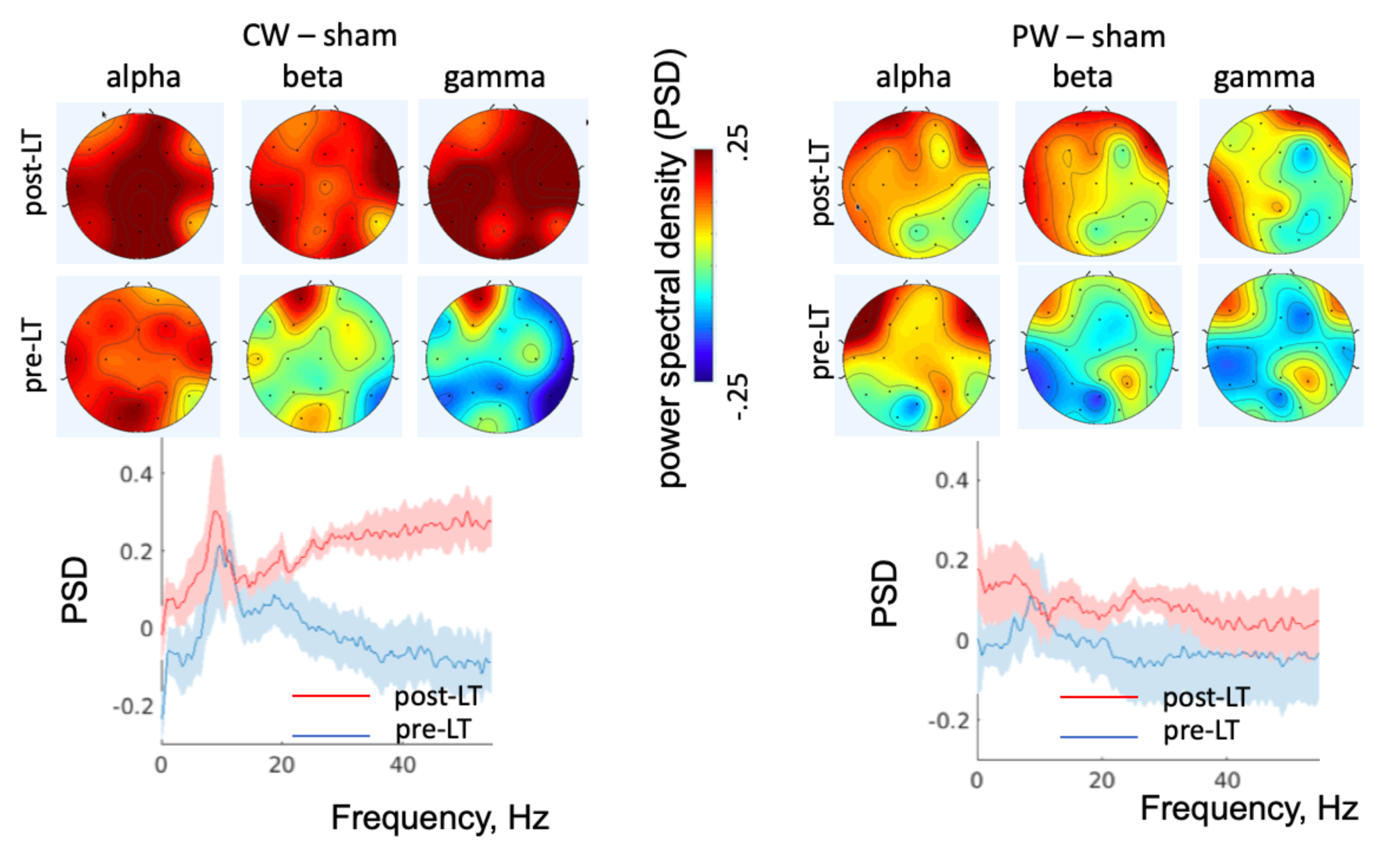
Eyes-closed resting state EEG: spectrogram differences before and after tPBM with continuous (CW) and pulse (PW) light (LT).

#### EEG Recordings During the Cognitive Task

Light-induced changes in EEG recorded while participants performed the 2-back task were similar to the changes described for the resting scans. We used a two-step approach to the analysis of EEG during the 2-back task. First (step 1), we directly compared EEG PSD obtained after each active treatment session to EEG PSD obtained after the sham session (always recorded during the 2-back task). In Figure 5, these differences (n = 5) are illustrated in the scalp maps on the top row, showing subtractions of post-c-tPBM (CW) PSD minus post-sham PSD in the left panel and subtractions of post-p-tPBM (PW) PSD minus post-sham PSD in the right panel. During 2-back performance, in the frontal-temporal scalp sites the post-c-tPBM PSD was increased relative to post-sham PSD. The global increase –averaged across all EEG channels– in post-c-tPBM PSD, relative to post-sham, was significant in the gamma band (mean change in γ-PSD_[POST (CW - Sham)]_ .1374 ± .0811 SD, t = 3.79, df = 4, p < .02). There was an effect trend in the beta band (mean change β-PSD_[POST (CW - Sham)]_ .1002± .0926 SD, t = 2.42, df = 4, p < .08). The effect was not significant for the post-p-tPBM comparison. Second (step 2), we examined if this difference between c-tPBM and sham visits was over and above any difference in the general state of the brain. We showed the post-c-tPBM vs. post-sham effect (recorded during the 2-back task) was larger than the pre-treatment difference in the PSD during the baseline resting scans with eyes open between these sessions (shown in bottom-row scalp maps in Figure 5). The global increase –averaged across all EEG channels– (n = 4 – in the bottom left panel) was larger in the post-c-tPBM vs. post-sham (during the 2-back task) relative to the pre-c-tPBM vs. pre-sham rest subtraction in the gamma (mean change in γ-PSD_[2-Back (CW - Sham) – PRE (CW - Sham)]_ .2811± .1755 SD, t = 3.20, df = 3, p < .05) and beta (mean change in β-PSD_[2-Back (CW - Sham) – PRE (CW - Sham)]_ .2256 ± .1049 SD, t = 4.30, df = 3, p < .03) bands. This difference was not observed for the p-tPBM condition. Despite these results are obtained in a small sample of participants, there is consistency in the c-tPBM treatment response across the subjects, which is reflected in statistical significance. Even though the n-back performance of participants suggests some practice effect as performance slightly improves from session 1 (CW) to session 2 (sham) to session 3 (PW), the increase in the brain activity (EEG) in the post-c-tPBM relative to post-sham condition occurred at session 1 relative to session 2; no improvements were noticed in the post-p-tPBM (session 3) relative to post sham condition (session 2).

**Figure 5.**
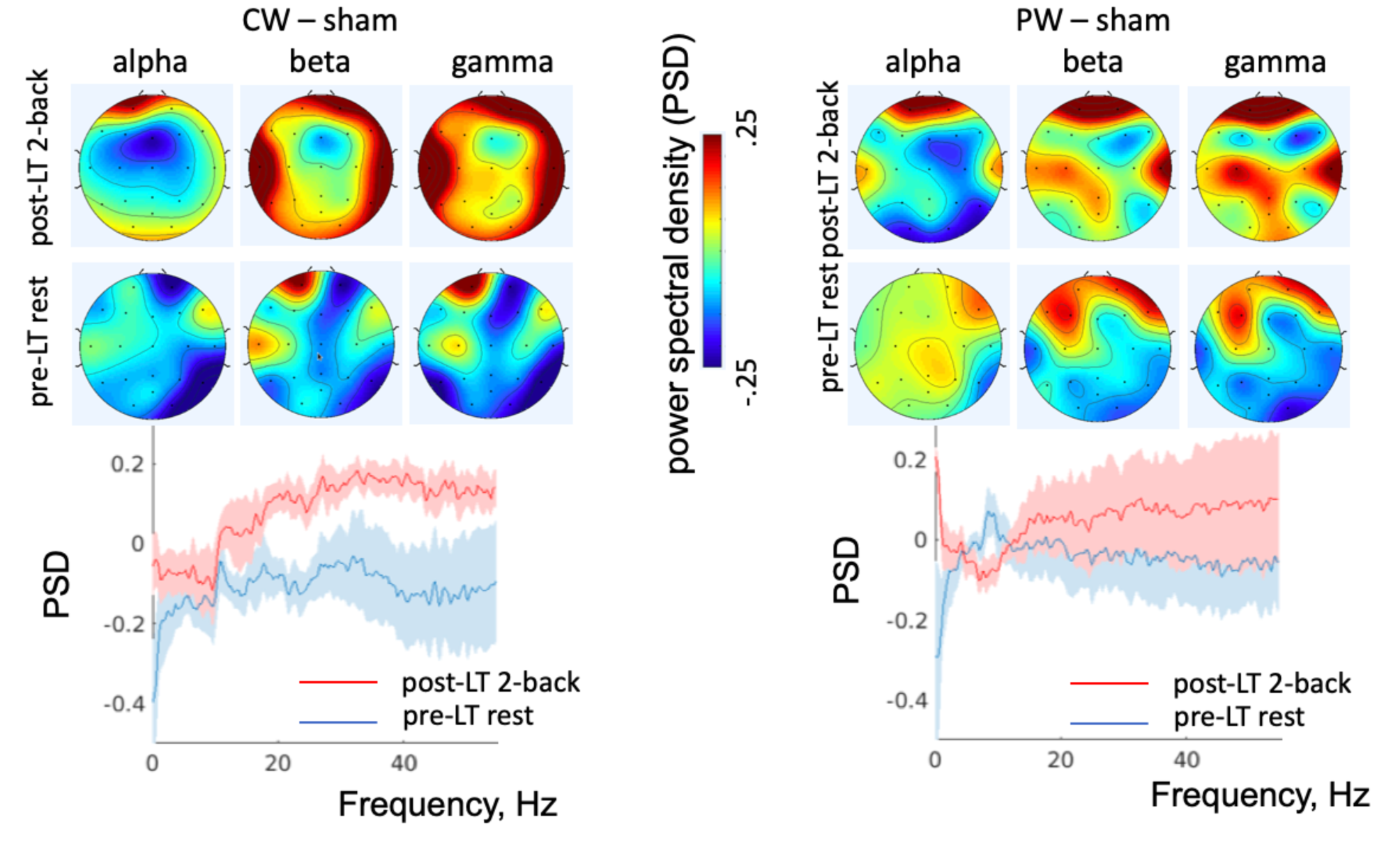
Eyes-open EEG during n-back task: spectrogram differences before tPBM (at rest) and after tPBM (during n-back), with continuous (CW) and pulse (PW) light (LT).

### 3.2 Cerebral blood flow (DCS)

In our participants’ sample (n=10), there were no statistically significant differences in the changes of DCS signals at each session (c-tPBM, s-tPBM, p-tPBM), for 2.5 cm and 5 mm separation and when subtracting the short separation to discount the contribution of superficial tissues to the measure of blood flow.

### 3.3 Cognitive Tests (n-Back)

We analyzed performance quality during the 2-back task. Figure 6 summarizes the levels of performance on each 2-back task at session 1 (n=6, c-tPBM/ CW). The performance was not significantly changed from pre- to post-c-tPBM treatment both in terms of accuracy in detecting targets (mean change −4.49 ± 20.84%, t = -.53, df = 6, p > .5) or reaction times [from 711.4316 ± 116.1718 SD to 733.9316 ± 61.1255 SD (sec), t = .45, df = 5, p > .5].

**Figure 6.**
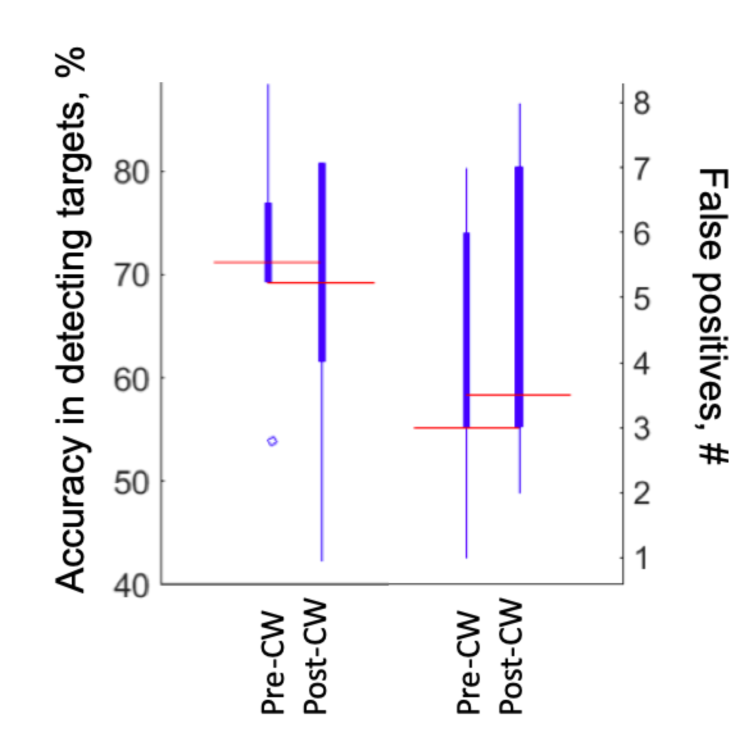
Performance on 2-back task: accuracy in target detection (%) and number of false postives (#) before and after tPBM with continuous (CW) light.

## 4. Discussion

Our study presents several important findings, worthy of being discussed: *1.* tPBM potentiated brain fast oscillations; *2.* tPBM-induced enhancement of brain fast oscillation was dose-dependent; *3.* tPBM enhancement of brain fast oscillation was unrelated to changes in cerebral blood flow (CBF), in fact tPBM did not affect CBF. These findings have broad implication for the field of neuromodulation with tPBM.

### tPBM potentiation of brain fast oscillations

Delivered by a four LED-clusters device in a single irradiation of 20 minutes to the forebrain, c-tPBM (CW) led to a significant increase in the high frequency neural activity, in the gamma and beta bands, believed to support higher-order cognition. During the resting state scans with eyes either open or closed, this increase was over broad frontal-temporal regions of the scalp, showing scalp topography common of the neural activity in these bands (Tewarie *et al*., 2016). During the 2-back working memory task, the increase reached significance in the beta band and was primarily over the frontal sites, likely because this task engages the prefrontal cortex (Spitzer & Haegens, 2017).

Our results are consistent with several previous studies of tPBM effects on the brain function. A recent investigation in healthy older adults at risk of cognitive decline reported an increase in the gamma EEG power, and a smaller increase in the beta power, over bilateral temporal scalp regions, during laser tPBM (CW, 1064 nm, 250 mW/ cm^2^, 137.5 J/cm^2^, 13.6 cm^2^ x 1 site) (Vargas *et al*., 2017). In the gamma band, this power enhancement persisted after tPBM. Another recent study with slightly different parameters of laser tPBM (CW, 1064 nm, 9.72 J/cm^2^ per minute, and 106.94 J/cm^2^ over 11 min), applied to the right forehead of healthy participants, showed an enhancement effect on the neural activity in high frequency bands (alpha and beta) (Wang *et al*., 2019). Whereas both these and the present study suggest a potential neuromodulatory effect, their differences in the analysis approach can explain some discrepancies in the findings. Our study which observed the tPBM effect on gamma and beta, not only compared the active tPBM to Sham (similarly to prior studies), but also measured the change from pre- to post-tPBM thus controlling for any additional differences between sessions (e.g., in the quality of EEG setup). The study by Vargas and colleagues took a within-session comparison approach without sham-control, which cannot partial out if participants were growing drowsier as they rested during tPBM. The study by Wang and colleagues did compare tPBM and Sham EEG measurements, however, measurements were collected on different days without accounting for potential between-session differences, such as in the EEG set-up, which gamma might be sensitive to.

Ability of tPBM to influence gamma band neural activity in the human brain may have important clinical significance. This type of brain activity has been linked to performance on complex and attention-demanding tasks (Tallon-Baudry, Kreiter, & Bertrand, 1999), and was implicated in support of diverse sensory and cognitive processes, including perceptual processing, object representations, visual awareness, and language (Senkowski & Herrmann, 2002) (Herrmann, Frund, & Lenz, 2010) (Tallon-Baudry & Bertrand, 1999) (Guntekin & Basar, 2014) (Busch, Herrmann, Muller, Lenz, & Gruber, 2006) (Busch, Debener, Kranczioch, Engel, & Herrmann, 2004) (Gruber, Tsivilis, Montaldi, & Muller, 2004) (Muller & Keil, 2004) (Keil, Muller, Ray, Gruber, & Elbert, 1999). Furthermore, it was found that the presentation of a previously learned stimulus evokes a stronger neural response in the gamma band then that of a new stimulus, suggesting a core role gamma activity may play in the mechanism of memory (Herrmann, Lenz, Junge, Busch, & Maess, 2004). Remarkably, clinical states, such as major depressive disorder, mild cognitive impairment, dementia due to Alzheimer’s disease and Down syndrome may alter gamma band brain activity (Basar, Basar-Eroglu, Guntekin, & Yener, 2013; Basar, Demiralp, Schurmann, Basar-Eroglu, & Ademoglu, 1999; Roh & Park, 2016).

### tPBM dose-dependent effect on fast brain oscillations

The effect of tPBM on high frequency brain oscillations disappeared when PW was used (p- tPBM). We interpreted this change as a dose-dependent effect, either due to the pulsed nature of PW tPBM (10 Hz, 33% duty cycle) –i.e., the on-off cycling of the light delivered affected the response– or due to the overall lower total energy delivered –0.8 kJ in PW tPBM compared to 2.3 KJ in CW tPBM–. A dose-dependent effect of laser tPBM with CW on behavior, accompanied by changes in the electrocorticogram spectra, was previously demonstrated in an animal model of depression (Mohammed, 2016). After pharmacological induction of depression, rats presented with both reduced survival behaviors on a forced swim test (FST) and reduced gamma-beta power at intracranial EEG. While laser tPBM led to normalization of behavior and electrocorticogram, only its lowest dose was effective; the middle dose produced no behavioral changes and the highest worsening of depressive behaviors. Observing a dose-dependent effect of tPBM on the neural activity in humans has great scientific significance as this suggests a causal link. It will be important in future research to further establish if different doses of tPBM influence the degree of the evoked neural changes. Characterizing the dose-response curve also has clinical significance, enabling the delivery of effective tPBM treatments. Our data suggests tPBM thresholds of insufficient (=<0.8kJ) and likely effective (>=2.3kJ) total energy for a single session paradigm in healthy subjects.

### Lack of tPBM effect on Cognition

Large enhancement on the resting state gamma and beta power, which was consistent across participants, suggests promise of tPBM for treatment of cognitive deficits. Intriguingly, the resting state gamma power has previously been linked to language and cognition during early development and in clinical conditions such as schizophrenia. Prior studies have also showed the effects of a single session of tPBM on cognition. For instance, Barrett and Gonzalez-Lima (2013) found that measures of attention (psychomotor vigilance task (PVT)) and memory (delayed match to sample (DMS)) improved in response to tPBM. In our study, we employed the n-back working memory task, which like PVT and DMS depends on the function of the prefrontal cortex. Unfortunately, in this pilot study the sample size was small, which is a likely explanation for why we failed to observe improvement on this task due to tPBM. Our very low accuracy rate at 2-back (67-71%), in a sample of mostly high-functioning and healthy subjects, also suggests that subjects might have underperformed, therefore jeopardizing the reliability of the task.

### Lack of tPBM effect on Cerebral Blood Flow

It was surprising that our study found no significant changes in the cerebral blood flow, as indexed by DCS recorded at 5 mm and at 2.5 cm source-detector separation in response to tPBM, either CW and PW mode, compared to the sham mode. In fact, to this date, one of the most validated and replicated neurophysiological findings in humans treated with tPBM is the increase in CBF (Salgado *et al*., 2015) . Furthermore, DCS, the technology used for CBF detection, is a validated technique with numerous studies supporting its use in humans.

Interestingly, prior studies − demonstrating an effect of tPBM on CBF− typically used laser devices and higher irradiance in the order of 250 mW/cm^2^. While technical failures in DCS are possible, they are also unlikely since our DCS probe was in close proximity of the light source and since our team is very experienced and conducts multiple DCS studies. It is more plausible that low-irradiance LED devices might selectively exert an effect on brain oscillation devoid of the effect on CBF. If this finding were confirmed, tPBM would likely have different neurophysiological mechanisms in humans depending on parameters used, such as light intensity or type of light source. If that were the case, tPBM could be better conceptualized as several treatment modalities, whose clinical efficacy and tolerability should be individually tested, rather than assuming a group effect.

### Strengths and Limitations

Several limitations should be acknowledged for our study: 1. This pilot study is based on a small sample of healthy participants; therefore, despite the strong effect on brain oscillations, the current results can only be considered preliminary. It is unproven whether the effects on brain oscillation would generalize to a wider population, including patients suffering brain disorders. 2. Similarly, due to the small study sample, we cannot judge conclusively the potential effects of low dose p-PBM (PW) on brain oscillations; it is striking that large effects were observed for CW, suggesting that, at the very least, effect sizes of PW and CW dosages were quite different. 3. It is also noteworthy that we only used a 20-channel EEG recording; it may be necessary to record with higher EEG sensor density to adequately characterize tPBM effects on brain oscillations (e.g. the effect of the p-tPBM over the anterior scalp regions). 4. Because the dose of tPBM was much lower in pulse mode (PW), this prevented from testing the impact of the pulsing of the light on brain oscillations. In the future, studies might adequately compare CW and PW effects, by matching average irradiance and total energy per session. 5. An additional limitation, also related to the study device, is the use of four clusters of LED sources mounted on a rigid frame. The rigid frame prevented the repositioning of two sources (on F3, F4) below the hairline; it is therefore likely that the actual dose of NIR was less due to the shielding effect of hair in this young cohort. 6. Our measure of the effects on cognitive function appears to be affected by the subjects’ poor effort at the n-back task, in addition to the small sample size. The strengths of our approach are: 1. In our study, tPBM delivered with an LED device was well tolerated with no serious adverse events. 2. LED tPBM has several advantages over laser, such as low or no risk of retina injury, lower cost, and potential self-administration. These features are decisive in terms of broadening of the clinical use of tPBM, as they offer a considerable progress in safety and comfort, compared to prior studies using laser sources. 3. The multimodal neurophysiological testing applied in our study offered a rare opportunity to understand the effects of tPBM in humans. 4. The single-blind, sham-controlled design, with careful match of all visible and auditory outputs of the tPBM device between active and sham sessions contributes to rigorous testing of our hypotheses. Of note, only two investigators (PC and EB) were always aware of the exact mode of tPBM applied at each session (CW, sham, PW), while investigators involved with neurophysiological testing remained mostly blind.

## 5. Conclusion

We observed a significant and large enhancement of the power spectral density of neural activity in the gamma-band in response to CW (c-tPBM) with NIR. This result is consistent with our hypothesis that tPBM influences high-frequency synchronized brain activity in the gamma band. By modulating the brain gamma activity, linked to higher-order cognition, tPBM might have a promise as a procognitive therapy.

It is encouraging that a large and significant increase in EEG metrics was found at rest with eyes open and closed and during cognitive challenge, despite the small sample of participants.

## Acknowledgments

The authors acknowledge the help of Arianna Riccio for drawing the depiction of our Transcranial Photobiomodulation device (TPBM-1000), Diffuse Correlation Spectroscopy (DCS) and electroencephalogram (EEG) systems (Figure 2).

None of the funding sources had any involvement neither in the study design, in the collection, analysis, interpretation of study data [but in the writing of the report, Luis DeTaboada offered his comments to the final manuscript], nor in the decision to submit the article for publication.

## Declaration of interest

### Dr. P. Cassano

Dr. Cassano has received consultation fees from Janssen Research and Development and from Niraxx Light Therapeutics Inc. Dr. Cassano has received unrestricted funding from PhotoThera Inc. and then from LiteCure LLC. to conduct studies on transcranial photobiomodulation for the treatment of major depressive disorder and a study on healthy subjects. He has also received funding from Cerebral Sciences to conduct a study on transcranial photobiomodulation for generalized anxiety disorder. Dr. Cassano cofounded a company (Niraxx Light Therapeutics Inc) focused on the development of new modalities of treatment based on near-infrared light. Dr. Cassano has filed several patents related to the use of nearinfrared light in psychiatry.

### Dr. Tatiana Sitnikova

Dr. Sitnikova was compensated for 6% of her full professional effort for approximately a year from Dr. Cassano’s LiteCure LLC. grant.

### Dr. E. Bui

Dr Bui receives royalties from Springer for a textbook.

### Mr. L. De Taboada

Mr. DeTaboada is a stock holder at PThera LLC and the CTO of LiteCure LLC.

### Dr. M. Hamblin

Dr. Hamblin is on the following Scientific Advisory Boards: Transdermal Cap Inc, Cleveland, OH; BeWell Global Inc, Wan Chai, Hong Kong; Hologenix Inc. Santa Monica, CA; LumiThera Inc, Poulsbo, WA; Vielight, Toronto, Canada;

Bright Photomedicine, Sao Paulo, Brazil; Quantum Dynamics LLC, Cambridge, MA; Global Photon Inc, Bee Cave, TX;

Medical Coherence, Boston MA; NeuroThera, Newark DE; JOOVV Inc, Minneapolis-St. Paul MN; AIRx Medical, Pleasanton CA; FIR Industries, Inc. Ramsey, NJ; UVLRx Therapeutics, Oldsmar, FL; Ultralux UV Inc, Lansing MI

Illumiheal & Petthera, Shoreline, WA; MB Lasertherapy, Houston, TX; ARRC LED, San Clemente, CA; Varuna Biomedical Corp. Incline Village, NV; Niraxx Light Therapeutics, Inc, Boston, MA;

Dr Hamblin has been a consultant for Lexington Int, Boca Raton, FL; USHIO Corp, Japan; Merck KGaA, Darmstadt, Germany; Philips Electronics Nederland B.V.; Johnson & Johnson Inc, Philadelphia, PA; Sanofi-Aventis Deutschland GmbH, Frankfurt am Main, Germany; Dr Hamblin is a stockholder in Global Photon Inc, Bee Cave, TX; Mitonix, Newark, DE.

### Dr. MA Franceschini

Dr. MAF has patents on the DCS technology and she is a consultant with Niraxx Light Therapeutics.

All other authors have no conflict of interest to declare.

